# Genetic identification and reiterated captures suggests that the *Astyanax mexicanus* El Pachón cavefish population is closed and declining

**DOI:** 10.1101/2022.12.01.518679

**Authors:** Laurent Legendre, Julie Rode, Isabelle Germon, Marie Pavie, Carla Quiviger, Maxime Policarpo, Julien Leclercq, Stéphane Père, Julien Fumey, Carole Hyacinthe, Patricia Ornelas-García, Luis Espinasa, Sylvie Rétaux, Didier Casane

## Abstract

The size of *Astyanax mexicanus* blind cavefish populations of North-East Mexico is a demographic parameter of great importance for investigating a variety of ecological, evolutionary and conservation issues. However, very few estimates have been obtained. For these mobile animals living in an environment difficult to explore as a whole, methods based on capture-mark-recapture are appropriate, but the feasibility of such approach and the interpretation of the results depend on several assumptions that must be carefully examined. Here, we provide evidence that minimally invasive genetic identification from captures at different time intervals can give insights on cavefish population size dynamics as well as other important demographic parameters of interest. We also provide tools to calibrate sampling and genotyping efforts necessary to reach a given level of precision. Our results suggest that the El Pachón cave population is currently very small, of an order of magnitude of a few hundreds of individuals, and is distributed in a relatively isolated area. The probable decline in population size in the El Pachón cave since the last census in 1971 raises serious conservation issues.

## Introduction

The Mexican tetra, or *Astyanax mexicanus*, is an outstanding freshwater fish model to study evolution. This species exists under two forms represented by cave-adapted and surface dwelling populations that can interbreed despite striking differences in morphology, physiology and behaviour (Keene et al., 2016). Today, there are 34 described cave locations which host *Astyanax* cavefish populations (Elliott, 2018; Espinasa et al., 2018; Espinasa et al., 2020; Mitchell et al., 1977). Genetic evidence suggests a recent origin of the cave populations of a few thousand years (< 20,000 years) according to some authors (Fumey et al., 2018; Policarpo et al., 2021) or up to about 150,000 years according to others (Herman et al., 2018). Whatever the date when extant troglomorphic populations first settled in caves and how many independent events were involved, genetic divergence between surface and cave fish is very low (Avise and Selander, 1972; Bradic et al., 2012; Fumey et al., 2018; Herman et al., 2018). The previous enlisted characteristics make it possible to use genetic methods to search for loci, and eventually to identify mutations, involved in the evolution of cavefish phenotypes (Casane and Rétaux, 2016; Protas et al., 2006). However, the environmental and demographic context should also be considered to better understand the evolutionary mechanisms involved in the fixation of these mutations, that is, the relative roles of selection and genetic drift. For a given subterranean population, the main demographic parameters of interest are its size and the migration rates between this population and other subterranean and surface populations. These parameters can vary across the distribution area and can change through time. Estimates of population size, dispersal potential and level of hybridization with surface fish, and their variations, are also important for conservation purposes.

For these small, mobile and relatively numerous animals that cannot be counted directly over their whole distribution area, population size can be estimated using capture-mark-recapture (CMR) methods (Bailey, 1951). The principle is as follows: we capture, mark and release *a* animals out of a total population of size *x*. After the marked animals have freely mingled with the unmarked, we re-catch a random sample of size *n*, of which *r* are found to be marked. Bailey obtained the maximum likelihood estimate of *x* and its variance:

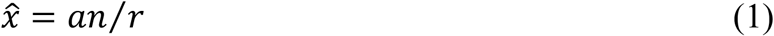

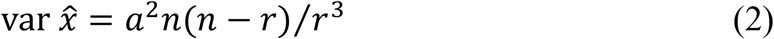

If *r* = 0 (no recapture of marked individuals), the population size cannot be estimated as 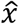 has an infinite expectation. If *r* is very small (a few marked individuals are recaptured), the variance is very large and we can get the order of magnitude of the population size rather than a precise estimate.

Excluding the case *r* = 0, Bailey found that the expectation of 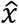 is biased, 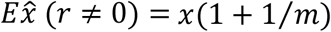, where m is the expectation of r. In order to take into account this bias when m is small, he proposed an adjusted estimate and gave its variance:

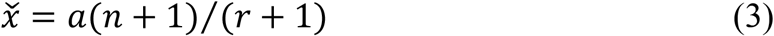

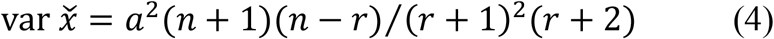

Equation (3) should not be used as a mathematical trick to solve the issue that a population size cannot be estimated if no marked animal is recaptured. Often, equation (3) and (4) are used instead of equations (1) and (2), not for obtaining precise and less biased estimates, but for getting an estimate with a smaller variance. However, the only valid solution to have a precise estimate, that is with a small variance, is to capture a large fraction of the total population in order to recapture many marked individuals. Moreover, the validity of the estimate depends on several assumptions that should be verified: 1) the population is closed, 2) all animals are equally likely to be captured, 3) capture and marking do not affect catchability, 4) marks are not lost.

The last two assumptions are a matter of the methodology. Researchers should use permanent marks and marking should be as little invasive as possible to limit effect on catchability. Whether the studied population is closed and restricted in a well-defined distribution area in which there is neither immigration nor emigration is often an unsettled issue. This is particularly true for cave animals for which we often can explore only a small part of their subterranean world. As marked and unmarked animals must have freely mingled before recapture, the animals must be mobile enough and the interval between sampling must be long enough to allow mixture. The validity of these assumptions depends on the model of the distribution of cavefish. We can imagine several possibilities, the two extremes being called here the “Oasis” and “Sea” model, and an intermediate “Lake” model (**Figure 1**). In the case of *Astyanax* cavefish populations, one model can be the most appropriate for a given region in the distribution area, but another model for another region. Moreover, in a given region, the best model can change through time. Even if models are oversimplifications of the reality, they help deciphering what is plausible from what is unlikely in a given case.

**Figure 1.**
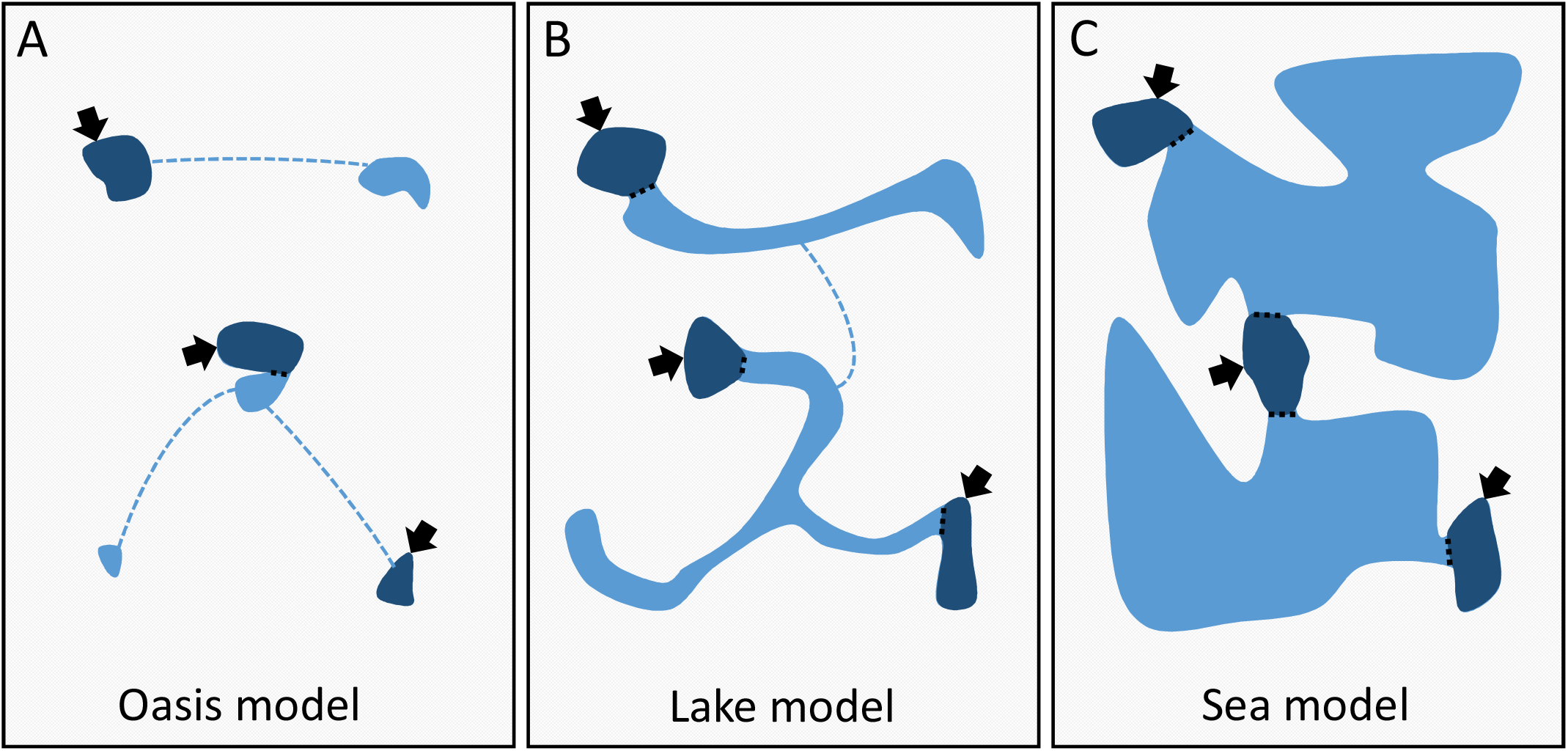
Hypothetical models for the distribution of cavefish populations. Dark blue represents water bodies that are accessible to humans and where fishes can be observed and sampled (arrows). Light blue represents large or restricted, permanent or temporary water conducts and bodies that connect accessible pieces of water. The “Oasis” model corresponds to a patchy distribution of small populations with little connections (dashed lines). At the other end, the “Sea” model corresponds to a large population of cavefishes distributed over a large area in fully connected aquifers. The “Lake” model is an intermediate. See text for details.

The “Oasis” model states that the fish distribution is very patchy, with few small isolated populations that are not permanently connected. It fits well the definition of closed populations. Some populations are accessible, so that population size and its variation through time can be estimated. In addition, if these populations have a relatively high extinction rate and if some empty areas are sporadically invaded, this system has the dynamic of a metapopulation. At the other extreme stands the “Sea” model. There, we have access to a few subterranean pools that are well connected to a single network over a large distribution area. In this case, the population of cavefish is large and homogeneous, and it is almost impossible to study its size using a CMR approach. The intermediate “Lake” model proposes the existence of clusters of caves, with caves well connected within a cluster, but clusters poorly interconnected. A cluster of caves fits the definition of a closed population, but a cave only gives access to a fraction of the population. Yet not impossible, it is difficult to study the demography of such large population. This discouraging outlook may have limited the research activity on cavefishes in this domain (Bichuette and Trajano, 2021; Elliott, 2018). Indeed, only two CMR studies have been performed on *Astyanax mexicanus* cavefish so far (Elliott, 2018). In March 1971, Elliot caught and marked 201 fish in Sótano de Yerbaniz and 230 in Cueva de El Pachón. A caudal fin clip was used to mark captured fish. One day and three days later, respectively, 226 and 242 fish were caught, among which 4 and 3 were marked. In El Pachón cave, he could see about 3 to 5 cavefish per m^2^, but the water was murky deeper than 60 - 90 cm, and he assumed there were more fish deeper. The visual estimate alone would have provided a minimal estimate of 950-1600 cavefish, but we now know that the full extent of the El Pachón pool is not visible to humans because the so-called Maryland extension is not accessible when water level is high. He kept 20 marked fish in a 19 L aquarium in the cave for the duration of the work to gauge the deleterious effects of the fin clip. In Yerbaniz, one control fish died and was eaten by the others after 48 hours. In El Pachón, five control fish died after 24 hours and one was nearly dead, but none showed obvious signs of attack by their fellows. Taking these death rates into account (1/20 and 6/20), the numbers of marked fish were corrected (191 and 161, respectively) and using equation (3) and (4), population size estimated to be 9,781 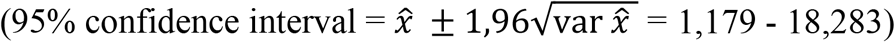 in El Pachón and 8,671 (1,810 - 15,534) in Yerbaniz. In 2009, Reynoso et al. captured and marked, by clipping the lower lobule of the caudal fin, 50 individuals in the El Pachón cave and 36 individuals in the Chica cave, but no marked fish were found among the 54 and 14 captured two days later, respectively, making it impossible to estimate the population sizes (Elliott, 2018). The absence of marked fish in the second sample in El Pachón and Chica could be the result of too small samples compared to the population sizes, but also the consequence of a high mortality of marked fish.

Undoubtedly, there is room to improve our knowledge of the demography of these subterranean fish as well as to develop novel non-invasive methods for CMR. Here, we examined the demography of the population in the El Pachón cave using a modified CMR method to better fit some assumptions and identify the most appropriate population model.

## Materials and Methods

### Fish sampling

Since 2004, we have maintained a laboratory stock of *Astyanax mexicanus* cavefish originating from the El Pachón cave (Sierra de El Abra, Mexico), initially obtained from W. R. Jeffery (University of Maryland, USA). In 2018 and 2022, 16 fish were sampled at random in our breeding facility for genotyping.

During field expeditions in the El Pachón cave, we sampled 20 individuals in 2016 and 35 individuals in 2019. In 2022, we sampled 30 and 29 individuals, respectively, at 3 days of interval (22nd and 25th February 2022), collected from the main pool.

In addition, in 2016, we sampled 38 surface fish individuals in the river Tampemole (thereafter called Arroyo Tampemole) and 8 surface fish individuals in a water extraction well close to the El Pachón cave (thereafter called Pozo Pachón Praxedis Guerrero) (**Figure 2**).

**Figure 2.**
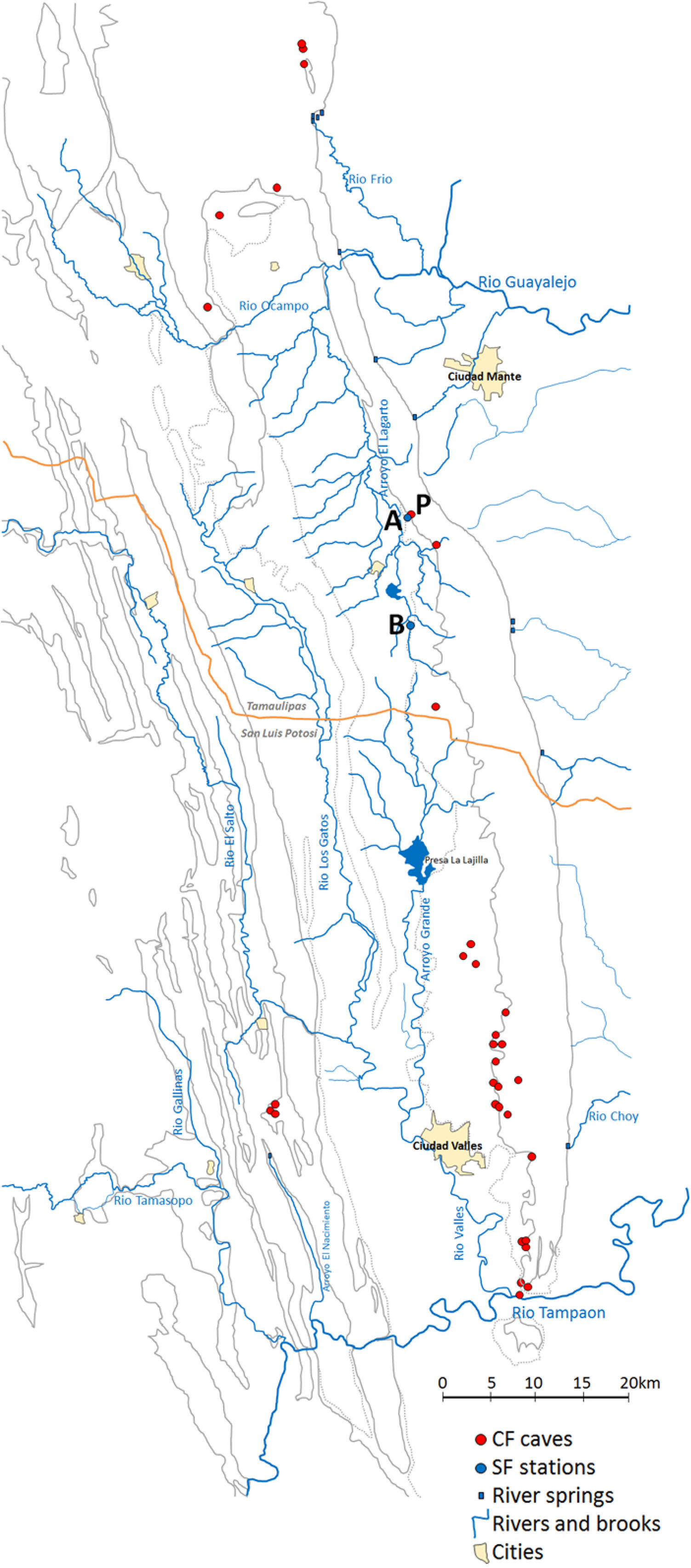
Map of the Sierra de El Ebra region in North East Mexico and sampling locations. Red dots indicate cave locations where cavefish populations have been described. P: El Pachón cave. Blue dots indicate sampling stations for surface fish, A: the well Pozo Pachón Praxedis Guerrero, B: the river Arroyo Tampemole.

### Fish genotyping

Up to 2022, all fish were fin-clipped and DNA extracted later from the fin fragments conserved in ethanol. Instead of this invasive fin clip procedure, which is the traditional way to get DNA, in 2022 we used a novel, non-invasive procedure. Gentle swabbings were performed on each flank of the fish in order to obtain two independent samples per individual. This procedure does not generate any physical damage to the fish and probably reduces stress significantly as compared to fin clipping. In all cases, fin-clipped or swabbed individuals were rapidly released in their natural pond after sampling.

The procedure of sample storage and DNA extraction was optimized to take into account the conditions of fieldwork, as follows.

DNA was extracted from swabs (FLOQSwabs^R^, COPAN Diagnostics Inc.) stored in tubes with silica gel beads at room temperature. Lysis with a mix of buffer (Tris HCl pH 8 100 mM, EDTA 2 mM, triton 0.2%) and proteinase K (250 μg/ml) overnight at 55°C was followed by inactivation of proteinase K for 10 min at 98°C. Aliquots of DNA were transferred in new tubes and stored at -20°C.

Among 26 microsatellite loci that proved to be highly polymorphic in this species (Bradic et al., 2012), we selected 18 loci on the basis that 1) they were polymorphic in El Pachón cave, 2) they could be easily amplified by PCR, 3) the amplicon sizes allow the amplification of these 18 loci through only three multiplexed PCR (**Supplementary Table S1**). Before setting up the PCR reactions, we prepared 10X primer mix with 2 µM of each primer (Multiplex1, Multiplex2 and Multiplex3). The PCR reactions were carried out in 10 µl of final volume with: 1 µl of template DNA, 5 µl 2X Platinum™ Multiplex PCR Master Mix (ThermoFisher Scientific), 1 µl 10X primer mix and 3 µl H_2_O. The program used was: 2 min 95°C, followed by 30 cycles of 30 sec 95°C, 90 sec 60°C, 60 sec 72°C), and a final extension for 30 min at 60°C.

Genotypes were scored using an ABI 3130 XL Genetic Analyzer, with GeneScan™ 500 LIZ™ size standard (ThermoFisher Scientific) and GeneMapper™ software v4.1 (Applied Biosystems™). For each locus, several alleles with different sizes were sequenced using homozygous specimens. These sequences allowed us to translate allele relative sizes obtained with GeneMapper into real allele sizes.

For each specimen captured in 2022, two independent samples (swabs) were genotyped. Globally, with most DNA samples (obtained by fin clip or swabbing, and from cave, river, well or lab individuals) we could obtain the genotype at each of the 18 loci, but for 8 fish the genotype at one or several loci was missing. These incomplete genotypes were not used in statistical analyses (**Supplementary Table S2**).

For statistical analyses, only complete genotypes were used. This included 16 samples from our breeding facility (2022); 19, 35, 29 and 29 samples from El Pachón cave collected in the field (2016, 2019, 22nd and 25th February 2022, respectively); 34 samples from Arroyo Tampemole, (2022), and 6 samples from Pozo Pachón Praxedis Guerrero (2022).

### Statistical analyses

#### Polymorphism variation through time

Changes in allele frequencies at each locus between each pair of sampling dates were tested using the Fisher’s exact test, using the fisher.test function from the stats package in R 4.1.0 (R Core Team, 2021). As 180 tests were performed, we used the Bonferroni correction (threshold α = 0.05/180) and Benjamini-Hochberg procedure in order to decrease the false discovery rate using the p.adjust function from the stats package in R (R Core Team, 2021).

#### Multidimensional scaling

To represent the matrix of genetic distances, the genetic distance between two individuals being the number of allelic differences, as a two-dimensional scatter plot, we performed a metric multidimensional scaling (MDS) using the cmdscale function from the stats package in R (R Core Team, 2021). We used only individuals for which a complete genotype was available.

#### Distribution of pairwise genetic distances according to kinship

First, we estimated the probability that two unrelated individuals would have the same genotype (*P*_*uni*_) using the following formula:

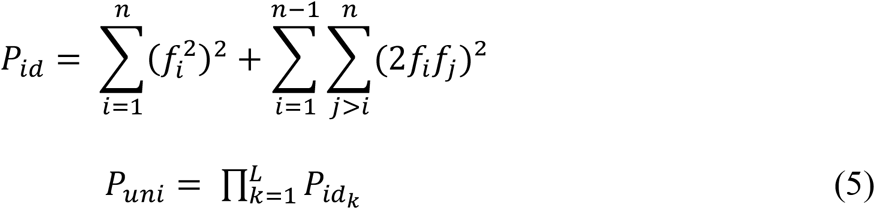

where P_*id*_ is the probability that two unrelated individuals have the same genotype at a given locus, *n* is the number of alleles at a given locus, *f*_*i*_ is the frequency of allele *i*, and *L* is the number of loci analyzed.

Although it is simple to compute the probability that two unrelated individuals have the same genotype, it is more difficult to compute the expected distribution of the genetic distance, that is the number of different alleles between two individuals, according to their kinship (*i*.*e*. unrelated, parent-descendant, full siblings, half siblings). These distributions were obtained using simulations. Pairs of genotypes were generated for different kinships. For two unrelated individuals, at each locus, the genotype of each individual was generated by randomly sampling two alleles, considering current allele frequencies in the population. For a parent and its descendant, the genotype of the parent was generated as described above, and the genotype of the descendant was generated by sampling at random one allele from this parent and the other allele from an unknown parent, that is from the population, considering allele frequencies. For full siblings, two unrelated parents were first generated; then, each descendant was generated by randomly sampling one allele from each parent. For half siblings, one mother and two unrelated fathers were first generated; then, one descendant was generated by randomly sampling one allele from the mother and one allele from one father. The other descendant was generated by randomly sampling one allele from the same mother and one allele from the other father. For each kind of relationship, one million simulations were performed to estimate the empirical distribution of pairwise distances according to a given kinship. Simulations and data analyses were carried out in the R statistical environment (R Core Team, 2021). The R script “GenerateIndividual_astyanax.r” can be found in GitHub “jmorode/Genetics_Astyanax”.

#### Estimation of genealogical relationships

The software ML Relate (Kalinowski et al., 2006) was used to find evidence of relatedness between cavefish. The accuracy of inferred genealogical relationships by the software was evaluated using simulated families of individuals of known genotypes. One thousand families were generated using the R script described above. Each family was composed by a mother, two unrelated fathers, two full siblings and two pairs of half siblings. This family composition allowed us to test the four relationships assessed by ML Relate: parent-offspring, full siblings, half siblings and unrelated. The genotypes of the members of these 1,000 families were written in an input file for ML Relate. For each pair of individuals in each family, the known relationship was compared with the one inferred by ML Relate. In each family, 6 unrelated individuals, 1 pair of full siblings, 2 pairs of half siblings and 6 pairs of parent-offspring were expected. The percentage of known relationships found by ML Relate over the 1,000 families was interpreted as an estimation of the accuracy of this software when identifying genealogical relationships in our observed population of cavefish.

## Results

### Genetic polymorphism in the El Pachón cave and nearby surface locations

Individuals collected in the El Pachón cave from 2016 to 2022 were genotyped at 18 microsatellite loci. In addition, El Pachón individuals from our lab stock, derived from El Pachón cavefish collected before 2004 (probably in the 1990’s), and surface fish from two locations, a well and a river close to the El Pachón cave, were also genotyped (**Figure 2**). All genotypes are reported in **Supplementary Table S2**.

The number of alleles at each locus was smaller in the El Pachón cave samples than in the two surface samples (**Table 1**). In total, among the 18 microsatellite loci analysed, only 54 alleles were recovered from 113 El Pachón cavefish whereas 203 alleles were recovered from 43 surface fish. The mean heterozygosity in El Pachón cave (0.2) is about 4 times lower than in river-dwelling fish (0.66 and 0.80 in Pozo Pachón Praxedis Guerrero and Arroyo Tampemole respectively).

**Table 1.**
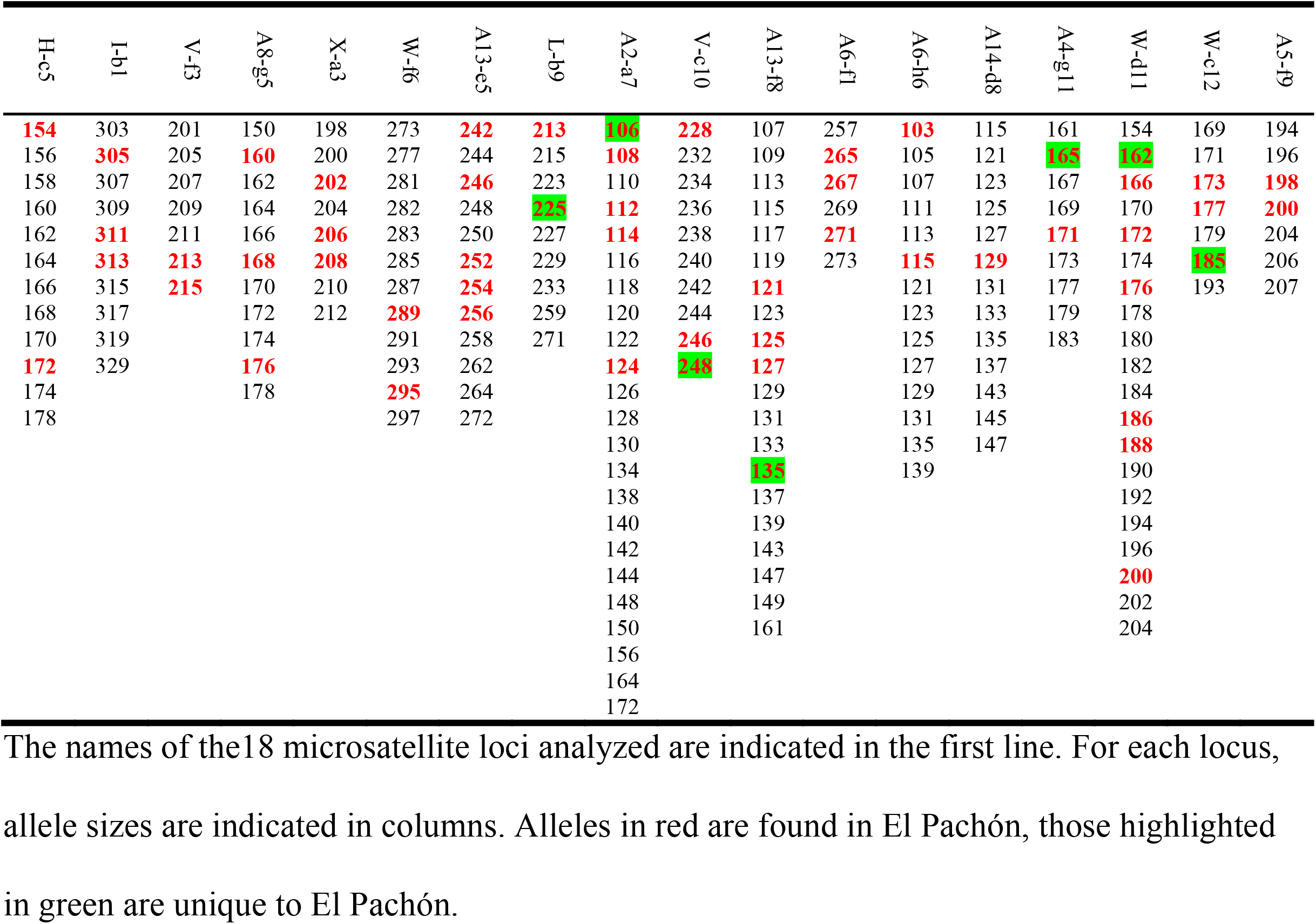
Allelic diversity in surface locations and El Pachón cave.

Allele frequencies are reported in **Table 2** for El Pachón cave samples and **Supplementary Table S3** for surface populations.

**Table 2.**
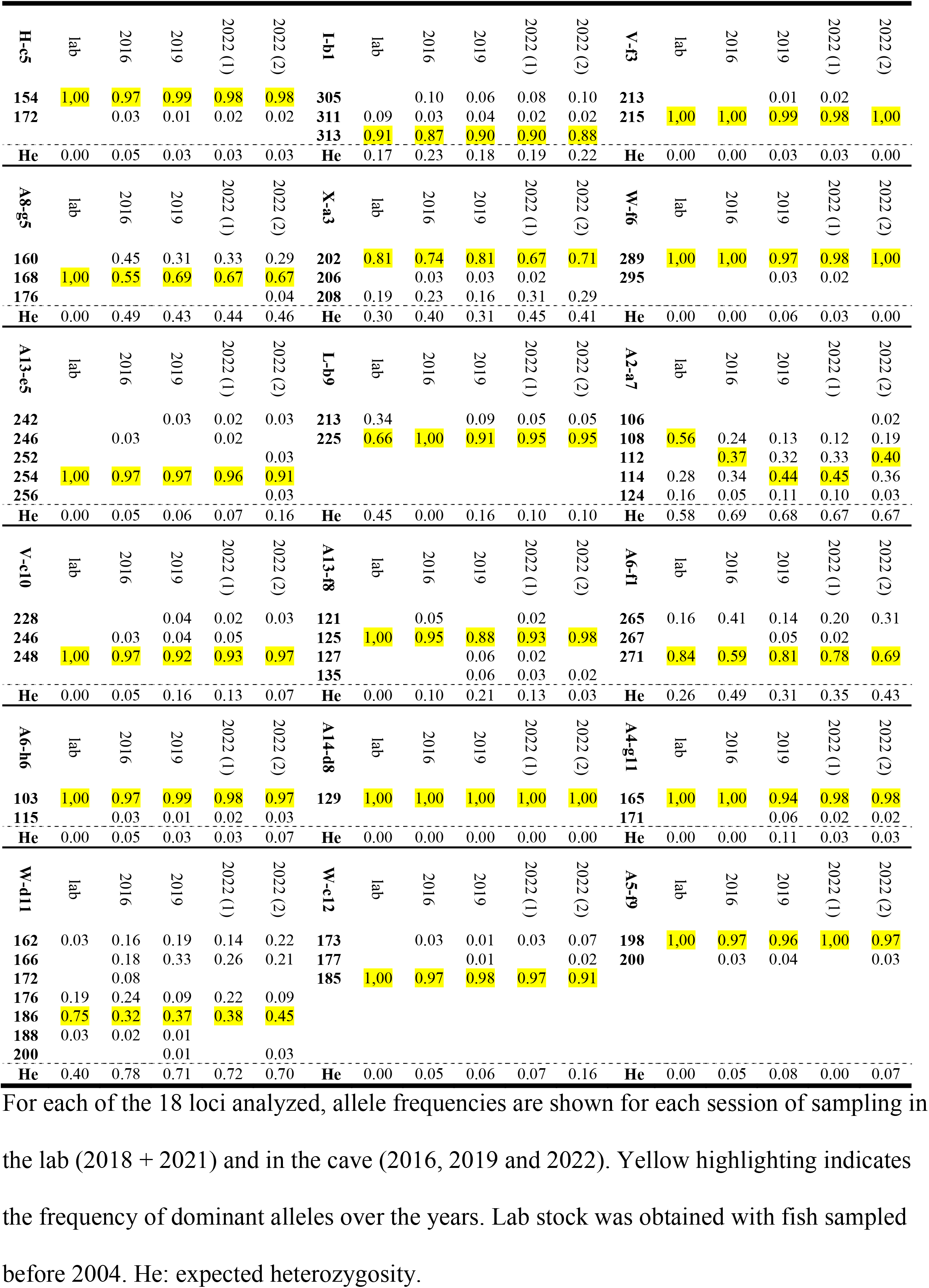
Allele frequencies in El Pachón cave.

The polymorphism in El Pachón cave appears stable since 2004, as all alleles recovered from lab individuals (i.e., derived from fish taken from the cave before 2004) are still present in the cave in 2022. In the lab stock, genetic drift may have led to the loss of some rare alleles (**Table 2, Figure 3**). Indeed, Fisher’s exact tests, with no correction for multiple tests (**Supplementary Figure S1A)**, with a Bonferroni correction (**Supplementary Figure S1B)** and applying the Benjamini-Hochberg procedure (**Supplementary Figure S1C)** indicate that significant allele frequencies differences observed between pairs of El Pachón samples involve most often the lab sample. However, these differences are minimal as compared to those observed between El Pachón samples and surface fish samples, or even between the two different surface fish locations (**Figure 3**, orange and green dots).

**Figure 3.**
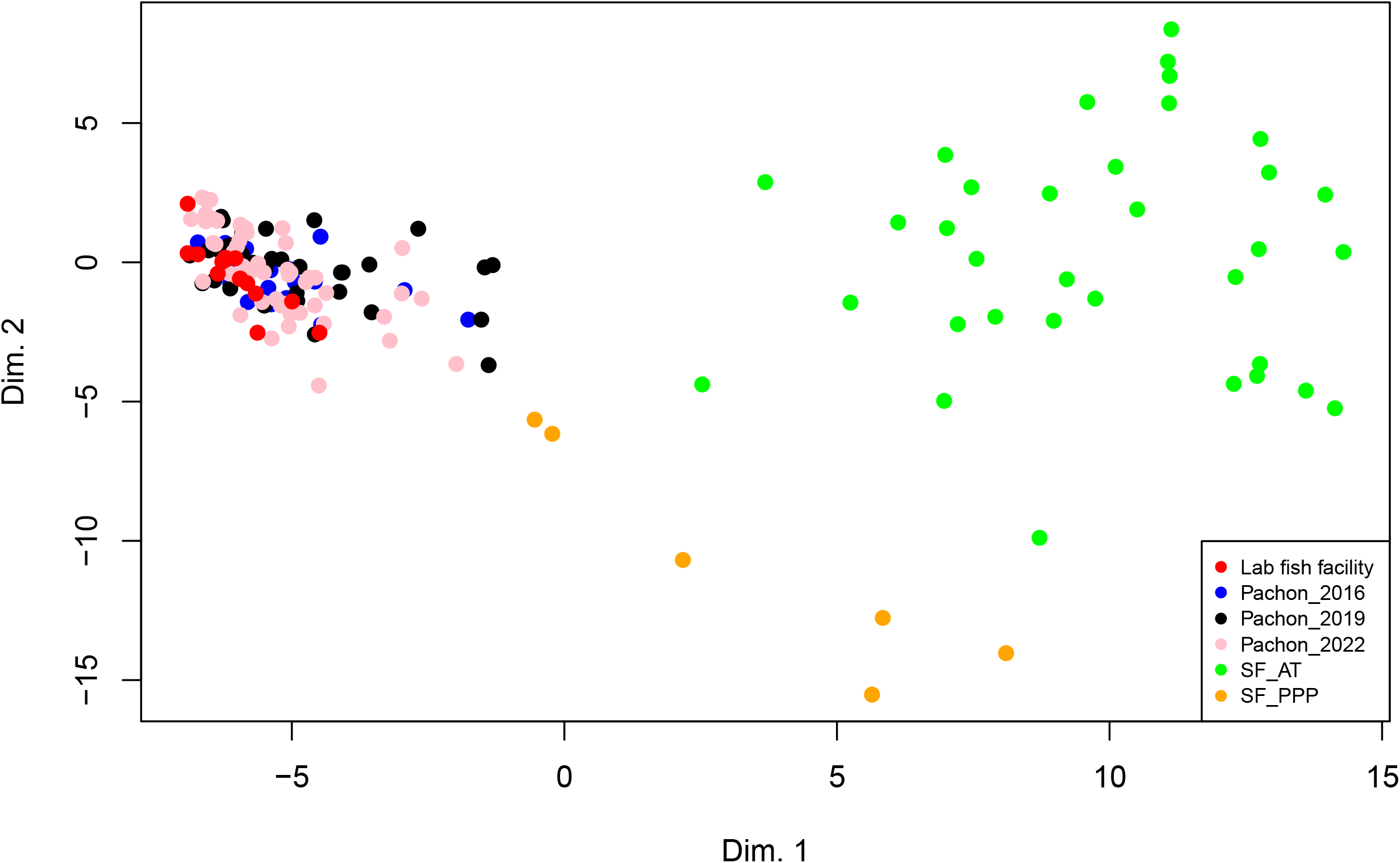
MultiDimensional Scaling (MDS) showing genetic distances between cave and surface fish. Each dot represents an individual. The color code for each sampling location is given in the inset. PPP = Pozo Pachón Praxedis Guerrero; AT = Arroyo Tampemole.

Of note, the polymorphism observed in our El Pachón cavefish samples is different from the one estimated in individuals collected in 2008 at the same place (Bradic et al., 2012) (**Supplementary Table S4**). The lengths of the alleles are different, and even doing a translation, we could not align the data. Moreover, higher polymorphism was found in Bradic et al. (2008). They often found two alleles differing by one repeat and with similar frequencies, while we find that for most loci one allele had a high frequency (> 80%). It is currently difficult to understand this discrepancy and these data were not further considered in our analyses.

### Genetic evidence of recaptures and population size estimation in El Pachón cave

The genetic capture-recapture method is based on the assumption that two fish have a very low probability of having the same genotype. The probability that, in our samples, two unrelated individuals have the same genotype was estimated using equation (5). For surface populations which display high allelic diversity, the probability is extremely low, 2.1 × 10^−15^ and 1.5 × 10^−23^ in Pozo Pachón Praxedis Guerrero and Arroyo Tampemole, respectively. For El Pachón cave, the probability is 2.1 × 10^−4^, 2.8 × 10^−4^, 2.2 × 10^−4^ and 2.2 × 10^−4^ for samples obtained in 2016, 2019 and 22 Feb 2022 and 25 Feb 2022, respectively. Thus, it is virtually impossible to catch unrelated surface fish with the same genotype, and it is unlikely for two unrelated cavefish to have the same genotype - despite the lower genetic polymorphism in the El Pachón cave population. One must also take into account that closely related fish, like full sibs, have more similar genotypes than unrelated individuals. Simulations of the genotypes of pairs of individuals with various relatedness were realized in order to estimate the probability of finding a given genetic distance, in the range of 0 to 36, according to kinship (**Supplementary Figure S2**). For full sibs (the most closely related individuals), in our simulations, we did not observe identical genotypes in surface populations, but about 2% in the El Pachón population. These results indicate that even if there were some closely related individuals in our samples of cavefish, the probability is low that they share the same genotype. In accordance with this assumption, we found four pairs of identical genotypes from different El Pachón samples, but none within a given sample. We therefore considered that identical genotypes in different samples corresponded to the same individual that had been recaptured.

On the 22nd February 2022, 29 fish were sampled by swabbing in El Pachón main pool and genotyped, and 29 fish were sampled 3 days later in the same place and genotyped. We identified 3 pairs of identical genotypes that we assumed to be recaptured individuals (**Supplementary Table S5**). Using equation (3) and (4), the estimated population size in El Pachón cave is therefore of 218 cavefish (95% CI = 40 - 395).

Interestingly, among 55 genotypes observed in 2022, we found three genotypes that were identical to genotypes observed in 2019, suggesting that three individuals have been recaptured three years after their first capture (**Supplementary Table S5**). Taking into account that 35 were captured in 2019 and assuming a low mortality between 2019 and 2022 of the fishes captured in 2019, using equation (3) and (4), we obtained another estimate of the population size in El Pachón cave, 490 (95% CI = 76 – 904). Both estimates point to a small population consisting of a few hundreds of individuals.

### Genealogical Relatedness

Individual genotypes from a small population give an opportunity to examine kinship. However, the reliability of kinship inferences depends on the level of polymorphism. We assessed the possibility to infer kinships among surface fish on the one hand and El Pachón cavefish on the other hand using the software ML Relate. First, pairs of individuals with different relatedness (unrelated individuals, offspring-parent, full sibs and half sibs) were simulated taking into account allele frequencies. Then the true kinship was compared to the one inferred by ML Relate. For surface populations, the accuracy was 81% and 92% for Pozo Pachon Praxedis Guerrero and Arroyo Tampemole respectively, suggesting that the polymorphism is sufficient to infer relatedness with a good confidence (**Supplementary Table S6**). However, for cavefish the accuracy was only about 50% (**Supplementary Table S6**). The polymorphism at the 18 microsatellite loci in El Pachón cave is thus not sufficient to identify related individuals.

## Discussion

### Genetic tags for long-term population surveys

Two assumptions about the experimental conditions, which should be verified for CMR analyses, are that capturing and marking have a minimal effect on the probability of recapturing. Several reports indicate that fin clip can affect their behaviour and be lethal for a substantial percentage of fish (Elliott, 2018). To circumvent this problem that is both technical and ethical, in 2022 we developed and used a method with swabbing instead of fin-clipping to collect DNA samples, which has been successfully applied for population estimates in amphibians including the flagship cave species, *Proteus anguinus* (Prunier et al., 2012; Trontelj and Zaksek, 2016). This approach is likely much less invasive if not completely neutral, if we do not take into account the stress induced by net capture and human handling during the swabbing procedure – which was performed as gently as possible. Yet it does “mark” captured fish genetically. Genotyping is then necessary to identify captured individuals. Although more time-consuming and costly, the great advantage of marking by genotyping is that we have access to a unique and stable tag that allows the identification of each animal over its entire life. However, the uniqueness of a genotype depends on the genetic diversity in the population. Microsatellite loci are particularly useful because their high mutation rate leads to several alleles at a given locus when the population size is relatively large, and individuals can be identified based on the combination of a small number of loci.

Conversely, in small populations, the polymorphism can be low and genotyping more loci might be necessary to associate a genotype with a unique individual. Here, we found that a combination of 18 microsatellite loci is sufficient to identify each El Pachón cavefish and compute population size, but the population is not polymorphic enough to infer kinship. The more reliable kinships inferred with surface fish polymorphism, which is about four times larger than in El Pachón cave, suggest that the use of four times more microsatellite markers would be necessary to get reliable inferences of kinship between cavefish.

### Size and isolation of the cavefish population in the El Pachón cave

There are four pools in El Pachón cave with sizable numbers of fish, but only two are usually accessible to humans: the main pool, which measures about 25 × 5 m and in which the study was conducted, and a small pool on a side lateral gallery about 3 × 1.5 m. These two pools connect during the rainy season. Fish are concentrated in the small pool at higher densities.

We have visited the El Pachón cave multiple times in the past 14 years and with photos and *in situ* observations, we estimate that in the small pool there are usually about 50-150 individuals. Often, a large part of the “main pool” is hidden to humans as water continues into the “Maryland extension” under a sump (Elliott, 2018). Only in 2003, 2009 and in February 2022 when we performed the genetic mark and recapture experiment have we seen the water level so extremely low as to give access to the Maryland extension galleries. In 2022, we visited the Maryland extension galleries and in particular the perched pool named UAEM pool (Elliott, 2018). This pool is higher and does not connect with the main pool. We did not do genetic mark and recapture experiments at the UAEM pool, but observations *in situ* suggest that the density of fish is similar to the main pool. The fourth pool was not visited in 2022, but in 2003 there may have been an estimate of about 50 individuals based on *in situ* observations. Given that during extreme rainy seasons all four pools may exchange individuals, the population size obtained for the main pool (218 fish) should be multiplied by ∼3 for El Pachón cave as a whole.

Our analysis therefore suggests that the El Pachón cavefish population size in 2022 was in the order of magnitude of a few hundreds. The capture in 2022 of three fish that were first captured in 2019 (with samples of 35 and 55 fishes, respectively) gives a population size of 490 fishes, which is in line with the estimate 654 (3 × 218) obtained with the CMR in 2022.

Together, these results suggest that the El Pachón population corresponds to the Oasis model, i.e., a small population poorly connected with other subterranean populations. Indeed, movements of fish between subterranean populations should prevent the recapture of fish over several years. Moreover, and importantly, the low genetic diversity and the stability of allele frequencies over time support the absence of a gene flow from surface or other genetically differentiated cave populations between ∼1990/2000 and 2022. In 1986 and 1988, some individuals with variable eye sizes and melanin pigmentation were observed in the albino and eye-reduced El Pachón population (Langecker et al., 1991). Thus, it is possible that sporadic migrations of surface fish occurred but the low and stable genetic polymorphism in the cave during the last 20 years when compared to the high polymorphism in close surface populations shows that surface fish migrations had an undetectable impact on the genetic diversity in the cave during the last few decades.

### Population genetic issues

Several researchers used genetic polymorphism to estimate the effective population size (N_e_) and the migration rate in El Pachón cave. The first one, examining the allozyme variations at 17 loci, found no polymorphism in this population, suggesting a small N_e_ and isolation from surface populations (Avise and Selander, 1972). Later on, the analysis of 26 microsatellite loci also pointed out a small N_e_ (< 1000) and limited gene flow (Bradic et al., 2012). A re-analysis of this data set, combined with transcriptomic data and using other statistical approaches led to the same conclusion (Fumey et al., 2018). Finally, a study using genomic data estimated that N_e_ could be higher (median = 32,000, min = 3,000 and max = 46,000) (Herman et al., 2018). N_e_ is a population genetic parameter that depends on the long term census population size (N_c_) but also on a series of other biological and demographic parameters (Charlesworth, 2009). In the case of *Astyanax* cavefish, two parameters could lead to a N_e_ much lower than N_c_, potentially several order of magnitude lower. First, episodes of low population size are known to have a disproportionate effect on the overall value of N_e_ (Charlesworth, 2009). Moreover, when females have the potential to lay thousands of eggs during their reproductive lifespan, like in this species, this can lead to a much larger variance in offspring number than expected with purely random variation, reducing N_e_ much below N_c_. Indeed, if many individuals from only a few egg laying events survive during exceptional and favorable environmental conditions whereas most individuals from most egg laying die, then the variance of reproductive success can be very large, a process known as sweepstakes reproductive success (Hedgecock, 1994). In a small population, this could result in many full- or half-siblings in a cohort sample. The hypothesis might be tested using genetic polymorphism. Even if the variance of reproductive success is not large, we can assume that N_e_ is at best in the order of magnitude of the smallest N_c_, that is a few hundreds.

### Conservation issue

In 1971, Elliott estimated the El Pachón cavefish population to ∼10.000 individuals (Elliott, 2018). Our 2022 estimate of the number of cavefishes in El Pachón cave to a few hundreds of fish, i.e., an order of magnitude lower, is very worrying, and questions the long-term maintenance of this population. The causes of the apparent decline in population size in this cave may be manifold. They might include variations in water quality parameters or water levels, human impacts like phreatic contamination, habitat disturbing like paintings of the cave walls above the main pool and plastic waste inside the caves, water pumping out of the cave for human consumption, and too frequent and too important samplings involving lethal experiments or removal of specimens from the cave by researchers, which are all facilitated by the ease of access to this cave. The effect by researchers should be easily and collectively handled and corrected by the *Astyanax* cavefish research community. Given all the intellectual benefit it has received from this single population, scientists should be actively and collectively engaged in its conservation. A longitudinal monitoring of the El Pachón cave population and the ecological parameters in the cave will also be paramount to counteract population decline and avoid extinction. Unfortunately and sadly, the El Pachón cavefish population, which has been the most studied and emblematic since its first scientific description in 1946 as “*Astyanax antrobius*” (Elliott, 2018) could well be the victim of its own success.

## Supporting information

Table S6_MLRelate_reliability

Table S5_Pachon_fish_genetic_distances

Table S3_allelic_frequencies_surface_fish

Figure S1A - Fisher

Figure S1B - corr Bonferonni

Figure S1C - corr BH

Figure S2_pairwise_genetic_distances

Table S2_Genotypes

Table S1_primers_multiplexed_PCR

Table S4_distrib_microsat_Pachon_in_Bradic_2012

## Data Availability

The R script “GenerateIndividual_astyanax.r” can be found in GitHub “jmorode/Genetics_Astyanax”.

## Supplementary Material

Supplementary data are available at Zoological Research

## Acknowledgments

This work was supported by a CNRS MITI (Mission pour les Initiatives Transverses et Interdisciplinaires) grant “Expérimentation en Milieux Extrêmes » to SR and collaborative grants from Agence Nationale de la Recherche (BLINDTEST) and Institut Diversité Ecologie et Evolution du Vivant to S.R. and D.C.

## Ethics approval

Animals were treated according to the French and European regulations for handling of animals in research. SR’s authorization for use of animals in research including *Astyanax mexicanus* is 91-116 and the Paris Centre-Sud Ethic Committee protocol authorization number related to this work is 2012–0052.

## Sampling authorization

*Astyanax mexicanus*: fin clips from surface and cave morphs of *A. mexicanus* were sampled during field expeditions, under the auspices of the field permits SGPA/DGVS/02438/16, SGPA/DGVS/1893/19 and SGPA/DGVS/03334/22 delivered to P Ornelas-García and S Rétaux by the Secretaría de Medio Ambiente y Recursos Naturales of Mexico (SEMARNAT).

## References

Avise JC & Selander RK. 1972. Evolutionary genetics of cave-dwelling fishes of genus Astyanax. Evolution, 26(1): 1–19.

Bailey NTJ. 1951. On Estimating the Size of Mobile Populations from Recapture Data. Biometrika, 38(3/4): 293–306.

Bichuette ME & Trajano E. 2021. Monitoring Brazilian Cavefish: Ecology and Conservation of Four Threatened Catfish of Genus Ituglanis (Siluriformes: Trichomycteridae) from Central Brazil. Diversity, 13(2): 91.

Bradic M, Beerli P, Garcia-De Leon FJ, Esquivel-Bobadilla S, Borowsky RL. 2012. Gene flow and population structure in the Mexican blind cavefish complex (Astyanax mexicanus). BMC Evol Biol, 12: 9.

Casane D & Rétaux S. 2016. Evolutionary Genetics of the Cavefish Astyanax mexicanus. In: Nicholas SF. Advances in Genetics. Academic Press, 117–159.

Charlesworth B. 2009. Effective population size and patterns of molecular evolution and variation. Nature Reviews Genetics, 10: 195.

Elliott W. 2018. The Astyanax caves of Mexico. Cavefishes of Tamaulipas, San Luis Potosi, and Guerrero. Association for Mexican Cave Studies. Bulletin 26: 1–325.

Espinasa L, Legendre L, Fumey J, Blin M, Rétaux S, Espinasa M. 2018. A new cave locality for Astyanax cavefish in Sierra de El Abra, Mexico. Subterranean Biology, 26: 39–53.

Espinasa L, Ornelas-García CP, Legendre L, Rétaux S, Best A, Gamboa-Miranda R, et al. 2020. Discovery of Two New Astyanax Cavefish Localities Leads to Further Understanding of the Species Biogeography. Diversity, 12(10): 368.

Fumey J, Hinaux H, Noirot C, Thermes C, Rétaux S, Casane D. 2018. Evidence for late Pleistocene origin of Astyanax mexicanus cavefish. Bmc Evolutionary Biology, 18(1): 43.

Hedgecock D. 1994. Does variance in reproductive success limit effective population sizes of marine organisms? In: Beaumont AR. Genetics and evolution of aquatic organisms. London: Chapman & Hall, 122–134.

Herman A, Brandvain Y, Weagley J, Jeffery WR, Keene AC, Kono TJY, et al. 2018. The role of gene flow in rapid and repeated evolution of cave-related traits in Mexican tetra, Astyanax mexicanus. Mol Ecol, 27(22): 4397–4416.

Kalinowski ST, Wagner AP, Taper ML. 2006. ML-RELATE: a computer program for maximum likelihood estimation of relatedness and relationship. Molecular Ecology Notes, 6(2): 576–579.

Keene AC, Yoshizawa M, Mcgaugh SE. 2016. Biology and evolution of the mexican cavefish. Academic Press.

Langecker TG, Wilkens H, Junge P. 1991. Introgressive hybridization in the Pachon Cave population of Astyanax fasciatus (Teleostei: Characidae). Ichthyological Exploration of Freshwaters, 2: 209–212.

Mitchell RW, Russell WH, Elliott WR. 1977. Mexican eyeless characin fishes, genus Astyanax: environment, distribution and evolution. Spec. Publ. Mus. Texas Techn. University, 12: 1–89.

Policarpo M, Fumey J, Lafargeas P, Naquin D, Thermes C, Naville M, et al. 2021. Contrasting gene decay in subterranean vertebrates: insights from cavefishes and fossorial mammals. Molecular Biology and Evolution, 38(2): 589–605.

Protas ME, Hersey C, Kochanek D, Zhou Y, Wilkens H, Jeffery WR, et al. 2006. Genetic analysis of cavefish reveals molecular convergence in the evolution of albinism. Nat Genet, 38(1): 107–111.

Prunier J, Kaufmann B, Grolet O, Picard D, Pompanon F, Joly P. 2012. Skin swabbing as a new efficient DNA sampling technique in amphibians, and 14 new microsatellite markers in the alpine newt (Ichthyosaura alpestris). Molecular Ecology Resources, 12(3): 524–531.

R Core Team. 2021. R: A Language and Environment for Statistical Computing. R Foundation for Statistical Computing, Vienna, Austria.

Trontelj P & Zaksek V. 2016. Genetic monitoring of Proteus populations. Natura Sloveniae, 18(1): 53–54.

