## Supplementary figures and images for "Genetic identification and reiterated captures suggests that the *Astyanax mexicanus* El Pachón cavefish population is closed and declining"

### Figure S1A - Fisher

Without any Correction

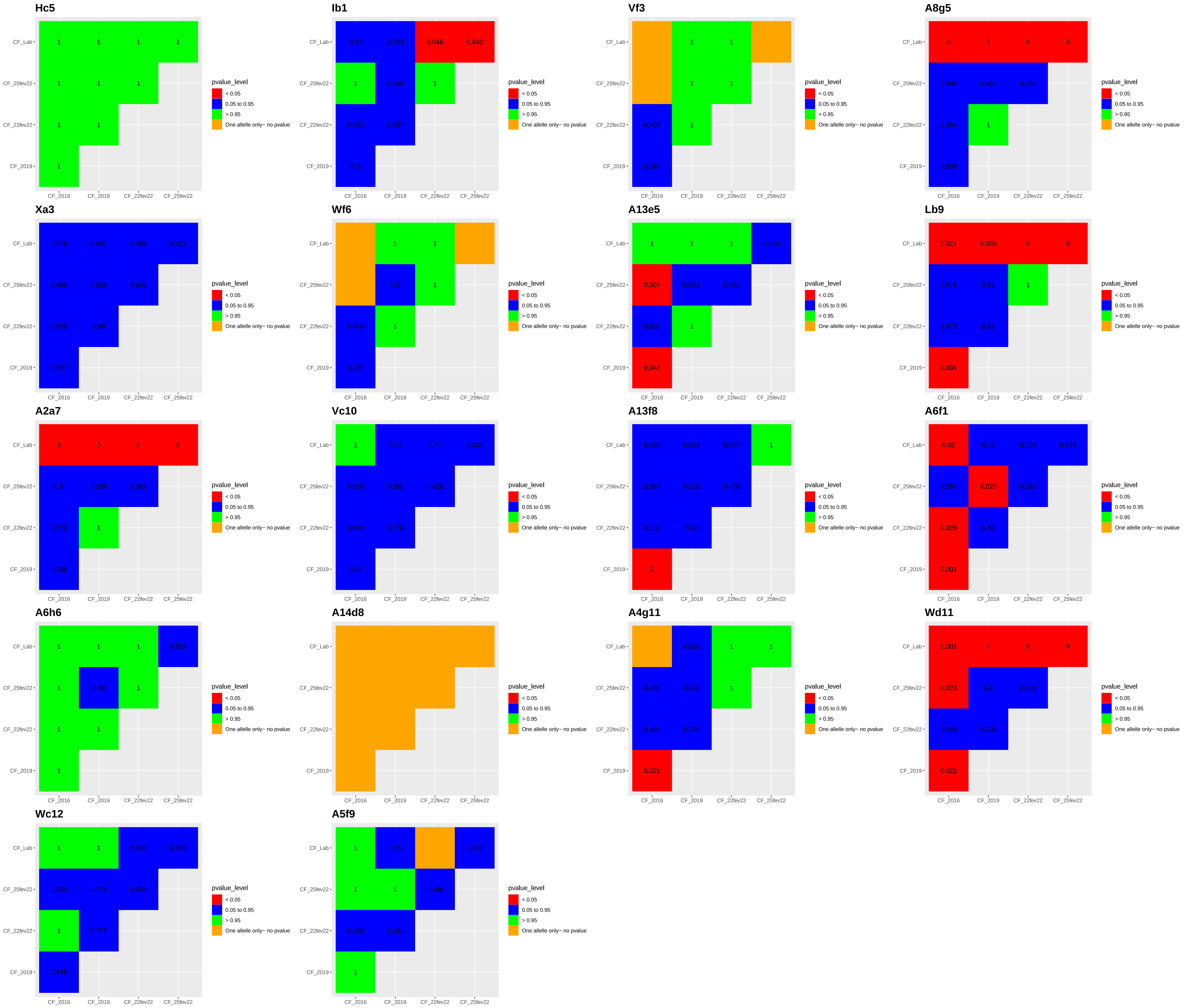

### Figure S1B - corr Bonferonni

Bonferroni Correction

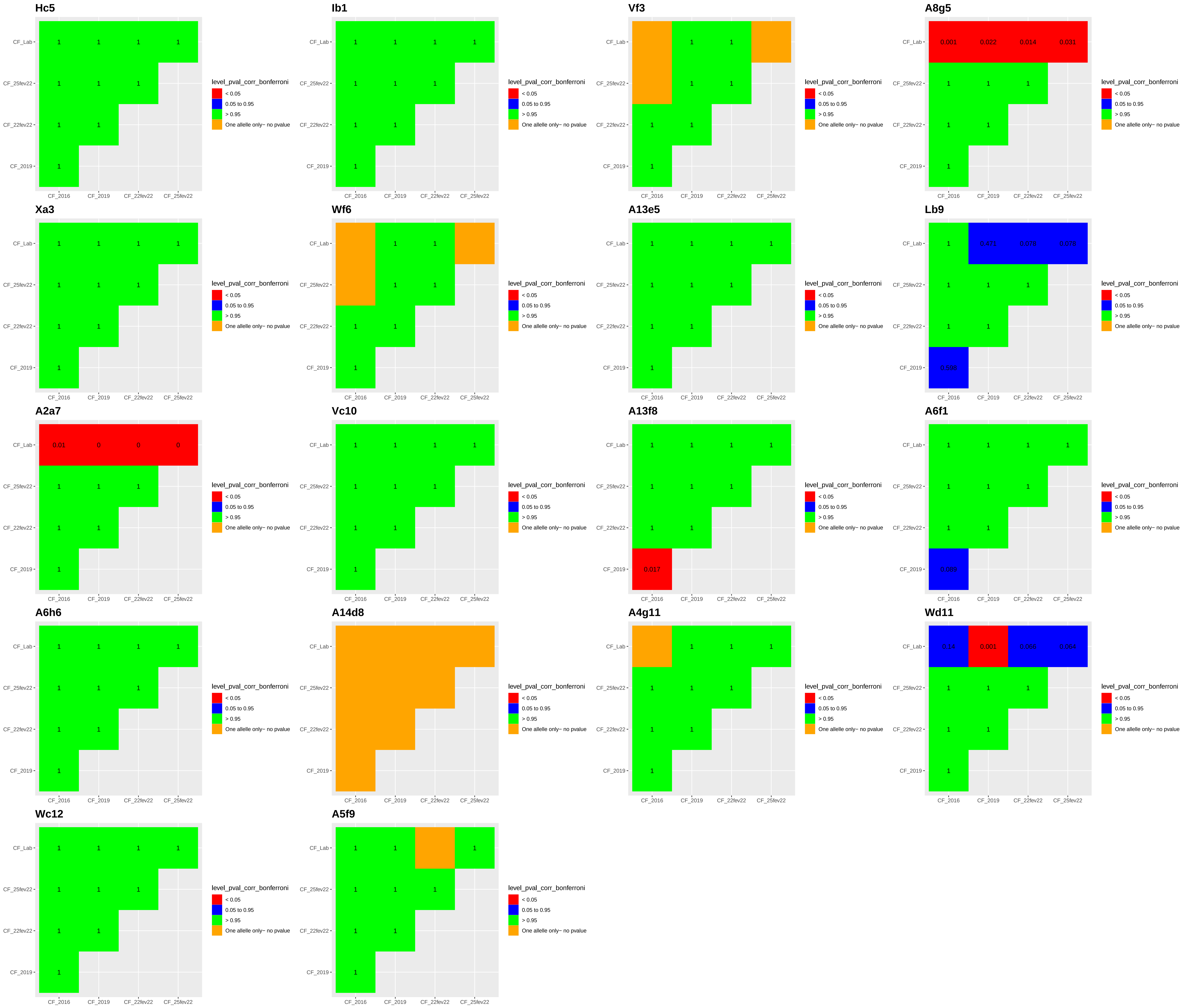

### Figure S1C - corr BH

Benjamini−Hochberg Correction

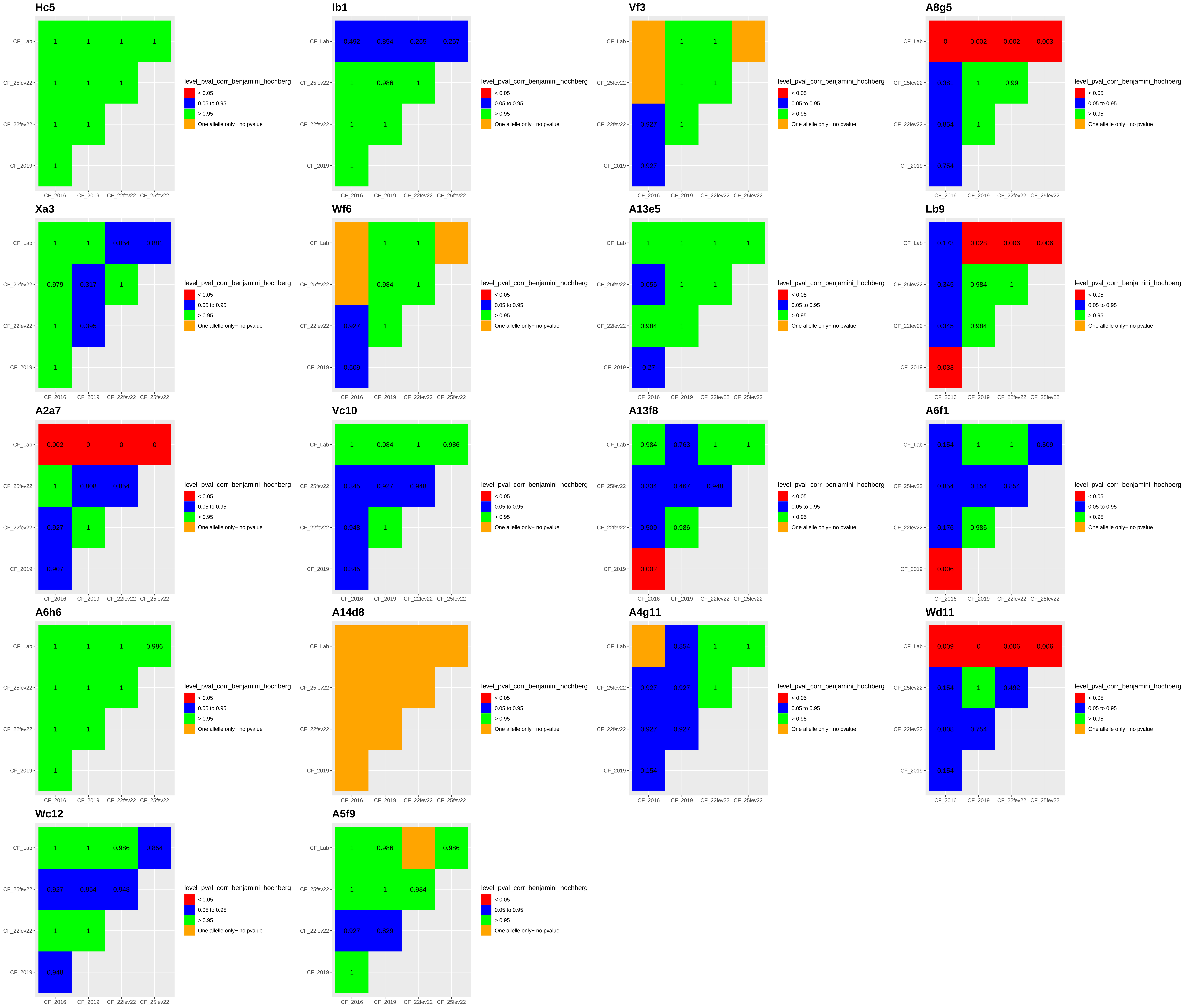
