## Supplementary material for "Genetic identification and reiterated captures suggests that the *Astyanax mexicanus* El Pachón cavefish population is closed and declining": Figure S2_pairwise_genetic_distances

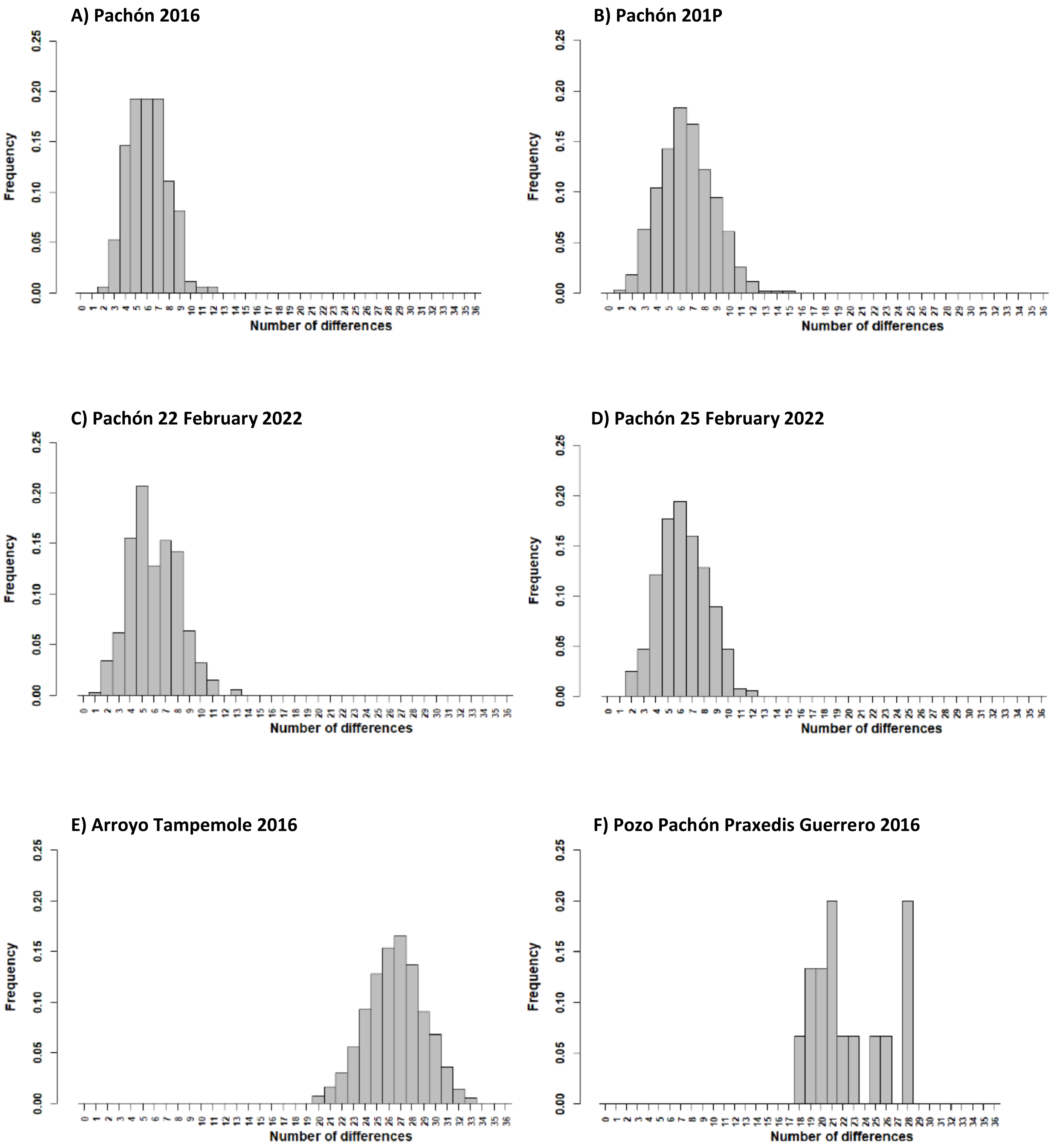
A) Pachón 2016
B) Pachón 201P
C) Pachón 22 February 2022
D) Pachón 25 February 2022
E) Arroyo Tampemole 2016
F) Pozo Pachón Praxedis Guerrero 2016
Frequency
Number of differences

G) Pachón 2016

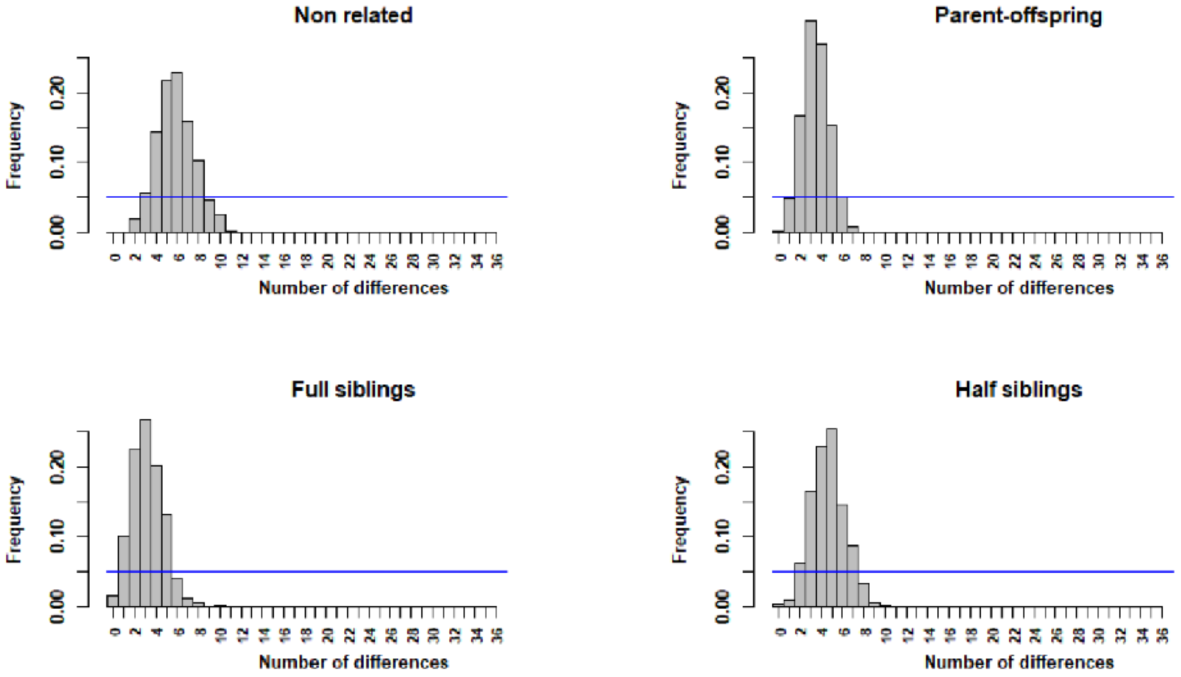


H) Pachón 2019

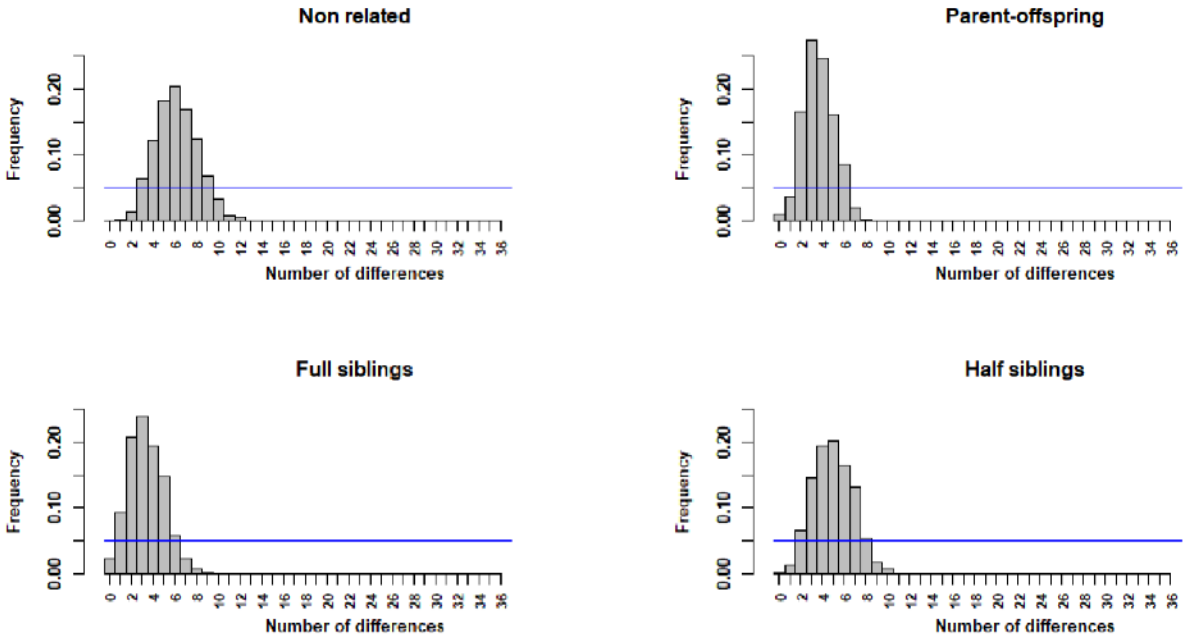

I) Pachón 22 February 2022

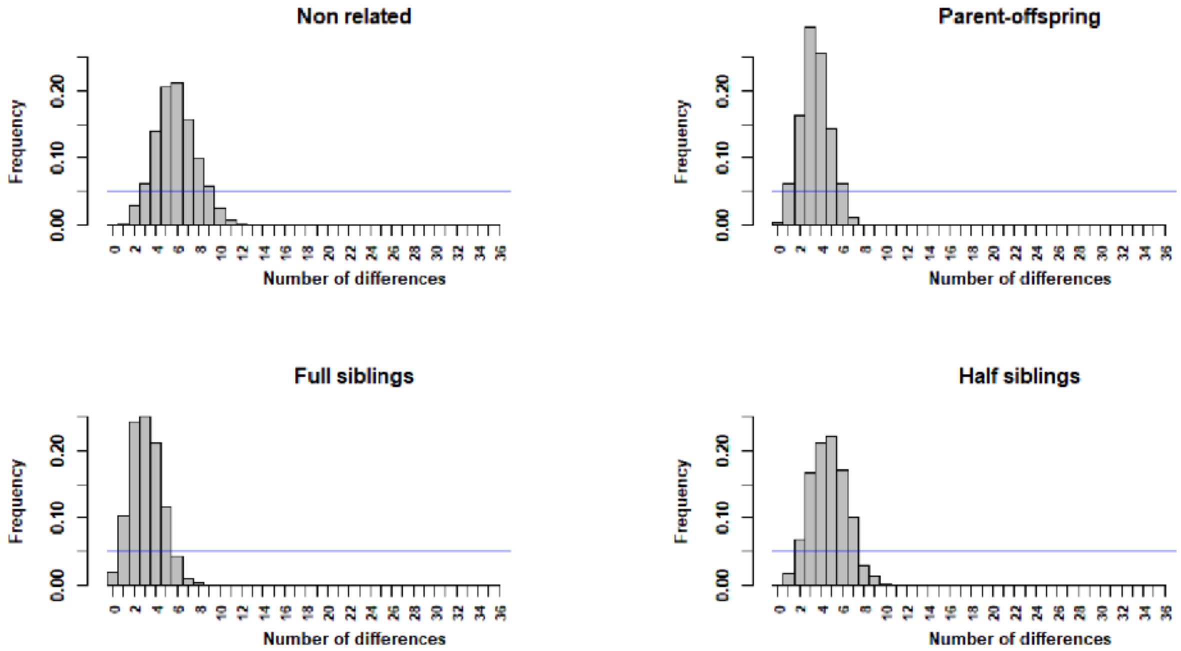


H) Pachón 25 February 2022

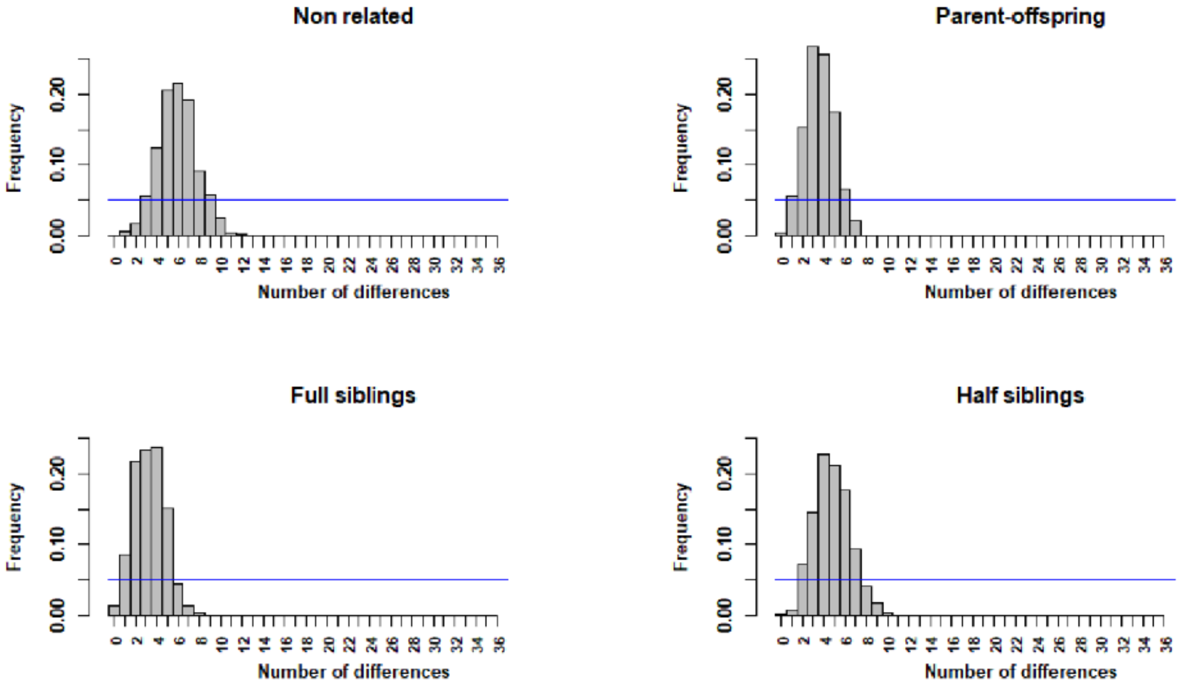

**K) Arroyo Tampemole 2016**

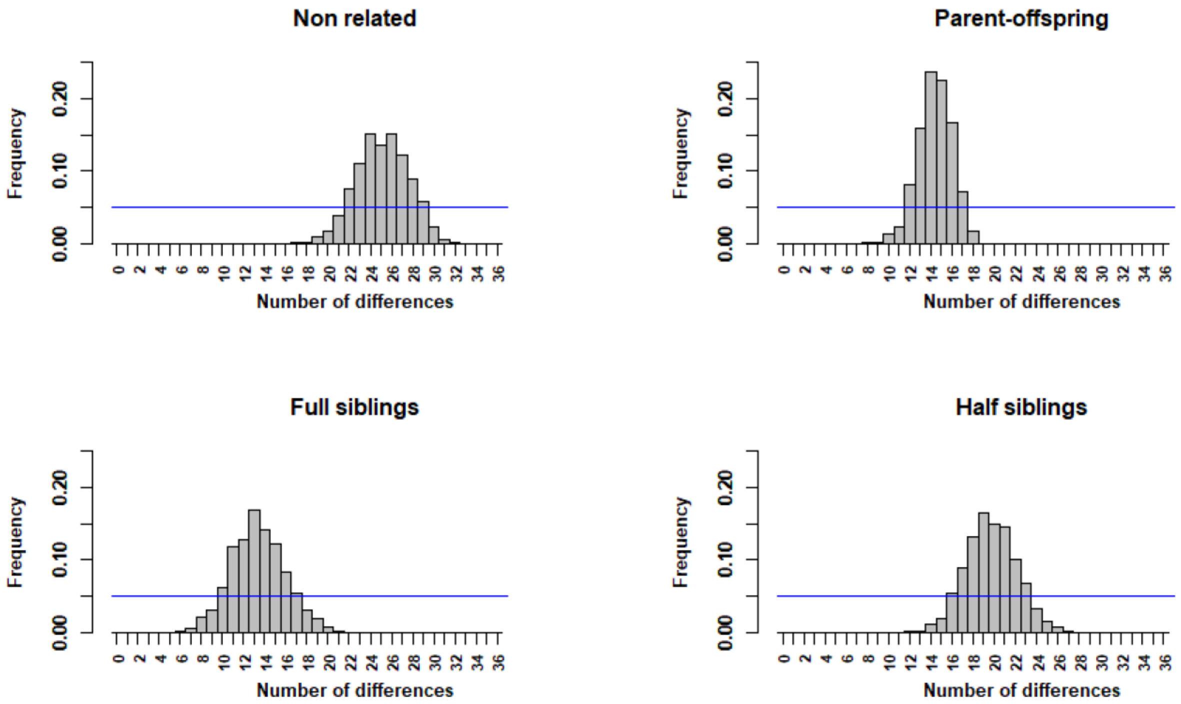


**L) Pozo Pachón Praxedis Guerrero 2016**

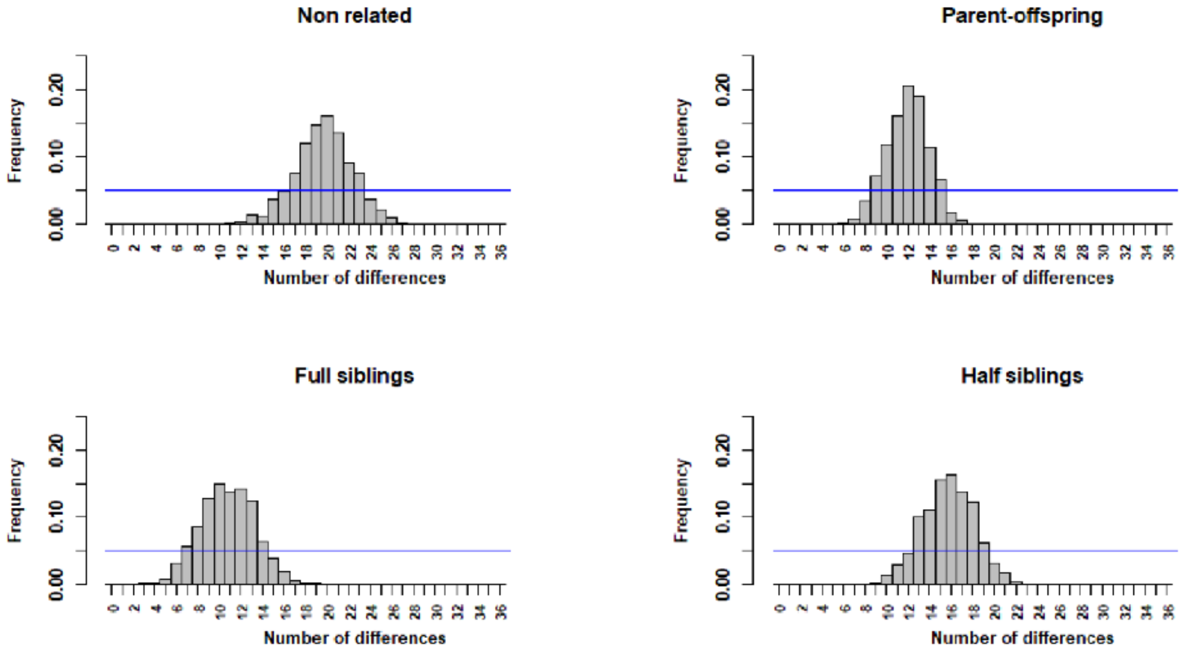
